# Subcellular resolution 3D light field imaging with genetically encoded voltage indicators

**DOI:** 10.1101/2020.05.22.108191

**Authors:** Peter Quicke, Carmel L. Howe, Pingfan Song, Herman Verinaz Jadan, Chenchen Song, Thomas Knöpfel, Mark Neil, Pier Luigi Dragotti, Simon R. Schultz, Amanda J. Foust

## Abstract

Light field microscopy (LFM) enables high signal-to-noise ratio (SNR), light efficient volume imaging at fast frame rates, and has been successfully applied to single-cell resolution functional neuronal calcium imaging. Voltage imaging with genetically encoded voltage indicators (GEVIs) stands to particularly benefit from light field microscopy’s volumetric imaging capability due to high required sampling rates, and limited probe brightness and functional sensitivity. Previous LFM studies have imaged GEVIs to track population-level interactions only in invertebrate preparations and without single cell resolution. Here we demonstrate sub-cellular resolution GEVI light field imaging in acute mouse brain slices resolving dendritic voltage signals localized in three dimensions. We characterize the effects of different light field reconstruction techniques on the SNR and signal localization and compare the SNR to fluorescence transients imaged in wide field. Our results demonstrate the potential of light field voltage imaging for studying dendritic integration and action potential propagation and backpropagation in 3 spatial dimensions.

## 1 Introduction

Cellular resolution voltage imaging enables direct observation of neuronal computation. Indeed, membrane potential imaging experiments have spatiotemporally resolved both active and passive action and synaptic potential generation throughout dendritic and axonal arbors.^1–14^ Resolution of these small voltage signals at high speeds requires high photon fluxes, making wide field single photon (1P) imaging by far the most common voltage imaging modality. Imaging neuronal processes with this technique requires the imaged membranes to lie approximately flat in the microscope’s focal plane. As these experiments are typically performed in slices, the requirement for flat, healthy, and superficial cells represents a significant barrier to entry for experimenters. Even in the best-prepared slices, anatomy dictates that only a few cells will be oriented parallel to the surface, reducing experimental throughput, and only certain cell types feature morphology that can be well-sampled by a single plane. Multiple approaches to improving wide field imaging’s 3D performance have been developed. Anselmi *et al*. (2011)^15^ applied remote focusing to axially shift and tilt the wide field focal plane as required by the sample, enabling calcium imaging along tilted dendrites. This adaptation, however, costs half of the fluorescence emission and is limited to a single tilted plane at a time. Point spread function (PSF) engineering via cubic phase masks^16^ or spherical aberration^17^ also enables parallelised volumetric sample imaging when combined with light sheet excitation, however to our knowledge these approaches have not successfully been implemented to image membrane voltage.

Lack of optical sectioning with wide field 1P imaging further complicates matters. Light from out-of-focus structures pollutes in-focus signals, confounding allocation of signals to axially separated processes. This issue is difficult to resolve with traditional optically sectioning confocal or two-photon microscopy approaches as they are point scanning. Sequential sampling of each pixel greatly reduces imaging bandwidth, and the fast frame rates required for voltage imaging necessitates short dwell times and therefore few collected photons. This restricts Poisson-noise limited SNR to low levels, making point scanning voltage imaging applicable to a limited number of experimental paradigms.^12, 18–20^

Fluorescence excitation parallelization into multiple spots,^21–27^ blobs,^28, 29^ lines,^30–32^ sheets,^33–40^ or specified patterns^41–45^ increases the photon budget, enabling functional volumetric imaging or single-plane imaging at increased speeds. A small number of these have been applied to imaging voltage in 2 dimensions,^26, 31, 42, 45^ however they are not able to image neuronal processes in 3D. Many of these techniques also trade-off reduced robustness to scattering compared to single-point scanning modalities for the increased excitation from parallelization.

Parallelized 3D two-photon imaging with elongated Bessel^46, 47^ or stereoscopic tilted^48^ beams excites narrow columns of fluorescence and relies on temporal and spatial sparsity of labelling and activity to demix time courses from different z planes. This increases the volume rate but each columnar pixel is still addressed sequentially, limiting bandwidth. These techniques have been used to image calcium fluorescence transients but not yet voltage.

Light field microscope (LFM)^49^ enables reconstruction of 3D volumes from single 2D camera images, extending wide field imaging whilst maintaining its unparalleled fluorescence excitation and collection efficiency. This is achieved by inserting a microlens array (MLA) at the native image plane of the microscope and placing the image sensor at its back focal plane (Fig. 1a). This disperses the angular components of the collected image (Fig. 1b), which can be used to infer objects’ axial positions. Each LFM image consists of circular subimages (Fig. 1c), with each subimage resembling a pixel in an undersampled image of the scene. Within each circular subimage, each pixel location encodes a different angular sampling through the object intersecting with the subimage’s location, a columnar tomographic projection through the sample.^50^ Light field images are typically parameterized by the 4D function *ℒ*(*u, v, x, y*), where each lenslet subimage is *ℒ*(*u, v, ·, ·*) and the same specific pixel under each subimage is *ℒ*(*·, ·, x, y*). The ‘native LFM resolution’ with which the object is laterally sampled is given by the microlens pitch divided by the objective magnification, much worse than the corresponding wide field resolution. In exchange the microlenses provide angular information that can be used to render views of the object from different perspectives, focus on different planes, and reconstruct 3D volumes all from a single 2D frame. This technique converts a key disadvantage of wide field 1P fluorescence excitation, lack of optical sectioning, into an advantage, as out-of-focus light renders 3D information about the sample.

**Fig 1.**
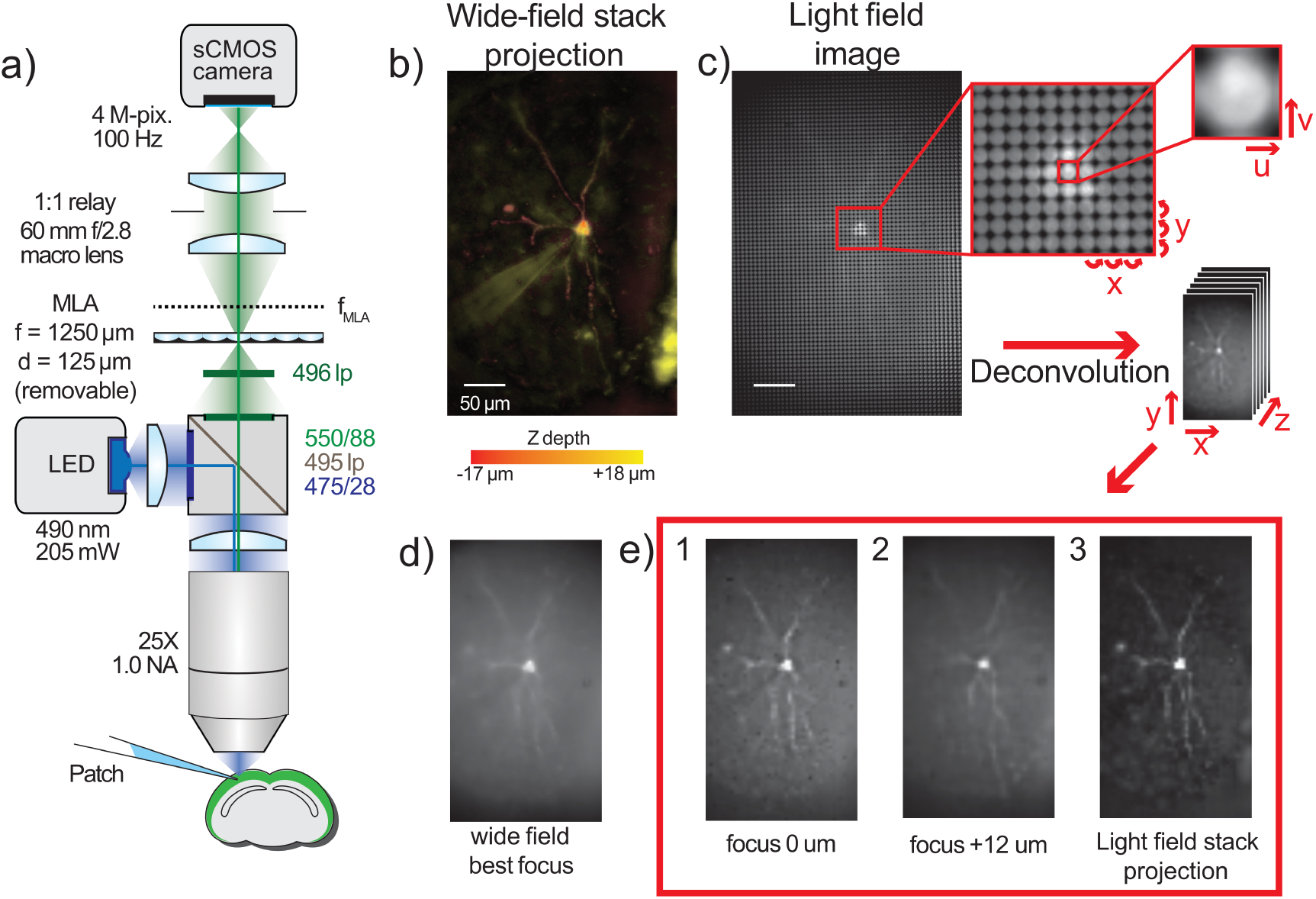
Light field microscopy enables simultaneous focusing on axially separated dendrites. a) LFM Diagram b) A pseudocolour z-projection of a wide field image stack through a GEVI labelled cell. Depth is color coded from red (superficial) to yellow (deep). Individual dendrites follow tortuous paths in all 3 dimensions, making simultaneous focussing on them all impossible in wide field microscopy. c) A light field image of the same cell showing the structure of light field images. Each spot in the light field image is a spatial sampling (coordinates x,y) of the angular distribution of rays (coordinates u,v) at that point. This angular and spatial information can be used to reconstruct a volume from a single image. d) A best focus wide field image of the single cell showing partially in focus dendritic structures. e) Three different images recovered from the light field image: 1 & 2) Single axial planes deconvolved showing individual dendrites seen out of focus in the wide field image. 3) A z-projection through the recovered light field volume image showing the in-focus sections of recovered dendrites. Figure adapted from Quicke (2019)^68^ CC BY-SA 4.0.

Two prominent algorithms for reconstructing source volumes from LFM images have been described, synthetic refocusing^49^ and 3D deconvolution.^50^ Synthetic refocusing relies on a ray optics model of LFM image formation to reconstruct images at the native LFM resolution equivalent to those of a wide field microscope focused at any axial plane in the sample. Focal stacks can be generated similar to standard microscope z-stacks by combining images reconstructed at multiple axial depths. Each pixel in the refocused image is a weighted sum of light field image pixels, meaning refocusing is fast. The reconstructed images, however, suffer from the same blur due to lack of optical sectioning as a standard wide field microscope.

An alternative approach is based on reconstruction of the source volume using a forward model of light-field image formation (the LFM PSF) based on wave optics.^50^ Iterative deconvolution approaches such as Richardson-Lucy (RL)^51, 52^ or the Image Space Reconstruction Algorithm (ISRA)^53^ find the maximum likelihood source volume given the measured image and LFM PSF in the presence of Poissonian (RL) or Gaussian (ISRA) noise. This approach is able to reconstruct source volumes at a lateral resolution greater than the native LFM resolution (the MLA pitch in the sample) by leveraging the fine sampling of the LFM tomographic projections.^50^ This increased resolution reconstruction fails where the tomographic sampling is degenerate, most notably around the native focal plane of the microscope, although newer designs have circumvented this limitation.^54–57^ The source volume is also reconstructed with less axial blur than in the refocused case, increasing axial signal discriminability.

Electrical length constants in neurons are on the scale of tens to hundreds of microns, making increased lateral pixel size less disadvantageous for voltage imaging. Over-resolving electrical fluctuations by imaging at or below the diffraction limit is typically unnecessary and can even hurt SNR by increasing the relative impact of non-Poisson noise such as read noise. Spatial resolution is therefore often sacrificed in voltage imaging experiments to increase speed or SNR. Many such experiments use low read noise, high sensitivity CCD sensors featuring low pixel counts, with pixels often measuring several microns across in the sample plane. Even with higher pixel count detectors, the relatively low sensitivity of many voltage probes means multiple pixels are often binned to increase SNR to acceptable levels. LFM’s decreased native lateral sampling rate therefore suits voltage imaging well, and deconvolution of LFM voltage imaging time series can be implemented without oversampling to reduce computational cost.

LFM has successfully imaged calcium over large volumes in C. elegans and zebrafish,^58, 59^ and in both head-fixed and behaving mice.^60–62^ Voltage dynamics have also been imaged successfully without single-cell resolution in Drosophila^63^ and larval zebrafish^64^ as part of whole brain imaging setups alongside calcium imaging. LFM has not, to our knowledge, been applied to studying subcellular or single-cell resolution voltage dynamics in any sample, despite its apparent suitability. In this study we apply LFM to sub-cellular GEVI imaging in acute mouse brain slices. We combine this technique with a recently reported transgenic strategy driving sparse expression in a random subset of layer 2/3 cortical pyramidal neurons which enables the resolution of single-cell level voltage signals in neuronal somata and dendrites.^65, 66^

We demonstrate that LFM is able to simultaneously image axially separated dendrites, enabling single-shot capture and localisation of GEVI fluorescence transients in the 3D dendritic arbour. We compare and evaluate deconvolution and synthetic refocusing for different GEVI imaging applications, whilst using a coarse deconvolution approach with no lateral oversampling to reduce computational cost. We also apply a recently developed LFM PSF calculation^67^ for high NA objectives. We show that LFM enables 3D localization of dendritic and somatic GEVI fluorescence transients and compare the extent to which refocused and deconvolved light fields enable lateral and axial transient localization. Finally, we compare temporal signal SNR between LFM and wide field microscopy.

## 2 Methods

This section reproduces methods described in Quicke (2019).^68^ We designed our LFM following the principles set out by Levoy *et al*. (2006).^49^ We adapted a wide field imaging system by placing a microlens array (MLA) at the microscope image plane, and used a 1:1 relay lens (Nikon 60 mm f/2.8 D AF Micro Nikkor Lens) system to image the MLA back focal plane onto our camera chip (ORCA Flash 4 V2, 2048 *×* 2048 pixels, 6.5 µm pixel size, Hamamatsu, see Figure 1a.). The lateral resolution is given by the MLA pitch divided by the magnification of the objective. Using our 25*×* objective (1.0 NA, XLPLN25XSVMP, Olympus) we chose our system to have 5 µm lateral pixels, dictating a microlens pitch of 125 µm.

The axial resolution is defined by the number of resolvable diffraction-limited spots behind each microlens.^49^ Assuming a central emission wavelength of 550 nm for mCitrine, the FRET donor in VSFP-Butterfly 1.2,^69^ the spot size in the camera plane is 6.46 µm using the Sparrow criterion. With an 125 µm pitch MLA, we are able to resolve *N*_*u*_ = 19 distinct spots under each microlens. The depth of field when synthetically refocussing our LFM can therefore be calculated as 7.81 µm.^49^

To efficiently use the camera sensor, the exit pupil of the objective should map through the MLA to produce circles on the light field plane that are just touching, requiring that the objective image-side f-number (f/12.5) equal the MLA f-number. We chose an f/10 MLA (MLA-S125-f10, RPC Photonics), an off-the-shelf part which came close to matching whilst being a larger aperture.

### 2.1 Imaging

This study was carried out in accordance with the recommendations of UK Animals (Scientific Procedures) Act 1986 under Home Office Project and Personal Licenses (project licenses 70/7818 and 70/9095). Slices were made from 4 mice aged 31, 32, 32 and 175 days transgenically modified to sparsely express VSFP Butterfly 1.2^69^ using the method previously described in Song *et al*. (2017).^65, 66^ These transgenic mice express the GEVI in cortical layer 2/3 pyramidal neurons under the intersectional control of TetO and destabilised Cre-recombinase.^70–72^ The destabilised Cre-recombinase was stochastically re-stabilised to induce sparse expression of the voltage indicator via two IP injections of a total of 2 *×* 10^*−*4^ mg kg^*−*1^ Trimethoprim (TMP, Sigma) over 2 consecutive days as described in Song *et al*. (2017).^66^

Slices were prepared at least 2 weeks post TMP injection using a method adapted from Ting et al. (2014)^73^ (the ‘protective recovery’ method, www.brainslicemethods.com). 400 µm slices were cut with a Camden Microtome 7000 in ice cold 95% O_2_ / 5% CO_2_ oxygenated ACSF containing: (in mM) 125 NaCl, 25 NaHCO_3_, 20 glucose 2.5 KCl, 1.25 NaH_2_PO_4_, 2 MgCl_2_, 2 CaCl_2_. The slices were then immediately transferred into NMDG-ACSF^73^ containing: (in mM) 110 N-methyl-D-Glucamine, 2.5 KCl, 1.2 NaH_2_PO_4_, 25 NaHCO_3_, 25 Glucose, 10 MgCl_2_, 0.5 CaCl_2_, adjusted to 300-310 mOsm/Kg, pH 7.3-7.4 with HCl and oxygenated with 95% O_2_ / 5% CO_2_ at 36 ^*°*^C for 12 minutes before being transferred back into the original sodium-containing ACSF for at least an hour before patching and imaging.

Fluorescent cells were patched under oblique infrared illumination (780 nm) with pipettes of resistances between 3 and 10 MOhms when filled with intracellular solution containing: (in mM) 130 K-Gluconate, 7 KCl, 4 ATP - Mg, 0.3 GTP - Na, 10 Phosphocreatine - Na, 10 HEPES. We digitized current clamp signals (Power 1401 digitizer; Cambridge Electronic Design) from a Multi-clamp 700B amplifier (Axon Instruments). At room temperature, we imaged at 100 frames/second for 2.5 seconds whilst injecting current pulses lasting 50 and 100 ms. Each pulse elicited depolarization to threshold evoking a single action potential or burst of 2-3 action potentials. We powered a 490 nm LED (M490L4, Thorlabs) with a constant current source (Keithley Sourcemeter 1401) to illuminate the sample at 3-11 mW/mm^2^. Sets of light field and wide field time series acquisitions were interleaved by removing and replacing the microlens array by hand. We averaged between 4 and 16 sweeps per imaging condition. The LED was collimated with an f = 16 mm aspheric lens (ACL25416U0-A, Thorlabs) and filtered with a 475/28 nm excitation filter (FITC-EX01-CLIN-25, Semrock). Fluorescence was collected using a 495 nm long pass dichroic (FF495-Di03, Semrock) along with a 550/88 nm collection filter (FF01-550/88, Semrock) and 496 long pass filter (Semrock FF01-496/LP) to attenuate any excitation light transmitted by the dichroic. Imaging data were acquired with Micromanager.^74^ Imaged cells’ somata lay between 11 and 40 µm below the slice surface, with a median depth of 29 µm. Data were analysed with custom Python scripts using SciPy packages.^75^

### 2.2 Light field reconstruction

We reconstructed source volumes using two techniques to compare their performance for single-cell voltage data. We calculated (x,y,z,t) volume time series using synthetic refocussing,^49^ and ISRA^53, 58^ using a PSF calculated using the method described in the section below. RL deconvolution^50–52^ was also tested on the data, however little discernible difference in the results was observed.

#### 2.2.1 Light field PSF calculation

We calculated LFM PSFs differently to previously described,^50^ using the method described in Quicke *et al*. (2019).^67, 68^ Briefly, to calculate the field at the microlens array we considered how a high NA objective lens collects the field from an oscillating electric dipole at position **r** near the microscope focus, |**r**| *<< f*, at the origin, calculating the Fourier transform of the field in the objective back focal plane. We assumed that we could model the behaviour of a point source consisting of randomly oriented fluorescent molecules as the incoherent sum of dipoles along 3 orthogonal directions. We then used the same method as described in Broxton *et al*. (2013)^50^ to model transmission through the MLA and to the camera.

We calculated the PSF for GEVI imaging deconvolution for 550 nm emission. We did not oversample the deconvolution as resolving voltage signals generally requires averaging pixels to approximately the native LFM resolution. We therefore generated a single light field kernel for each depth by averaging over kernels sampled for point sources at different lateral positions under the microlens, weighting each point in the average by a 2D Hamming window function of a width equal to our microlens’ pitch. We averaged over kernels sampled at 5 times finer than the native microlens resolution. The ISRA was used to deconvolve the data.

#### 2.2.2 Volume reconstructions

Having obtained our downsampled PSF we deconvolved our volume using a similar procedure to previous studies. A key difference is that only a single 2D convolution was required for each depth in the reconstructed volume for the forwards and backwards projections, respectively, as we did not increase the lateral sampling rate. We applied the deconvolution scheme independently to each frame of the image time sequences, using a cluster to parallelize the data processing. Deconvolution of a single frame took around 30 - 40 minutes for a 21 iteration deconvolution of 21 z-planes on a single CPU. We employed a large cluster to process the individual frames simultaneously, enabling 5000 frames to be processed overnight. We did not use a parallel algorithm within each deconvolution to leverage, e.g., GPU processing, as the computing resources available to us were better suited to data parallelism. As with previous studies this would greatly increase the rate of individual frames, although it would also likely reduce the number of simultaneous frames that could be deconvolved for typical cluster setups.

Synthetic refocusing, based on a ray optics model of light field image formation, is a simpler approach to volume reconstruction that is also much less computationally intensive. Images focused at different z-depths can be constructed by combining individual perspective views using the formula derived in Ng *et al*. (2005).^76^ Linear interpolation in this summation results in each pixel being the weighted sum of pixels of the original light field image. This reconstruction is much faster than the iterative deconvolution methods and also does not suffer from noise amplification.^77^

### 2.3 Volume time series analysis

#### 2.3.1 Effect of reconstruction on SNR

To compare the effect of different reconstruction techniques on voltage signal SNR we reconstructed single planes from volumes at the light field microscope focus. We compared synthetically refocused time series with time series deconvolved using ISRA for different iteration numbers. Regions of interest (ROIs) were manually chosen over the soma and its surround and were identical for the synthetically refocused and deconvolved volumes.

As we were collecting fluorescence from the VSFP Butterfly 1.2 FRET donor, fluorescence decreased upon membrane depolarisation.^69^ Therefore the traces shown Figures 2 and 3 are inverted. To measure SNR we calculated the signal as the 5^th^ percentile value during a stimulus and relaxation period of 200 ms with the median value of the 100 ms before the stimulation period subtracted. The noise level was calculated as the standard deviation of a 350 ms period during no intracellular current stimulus.

**Fig 2.**
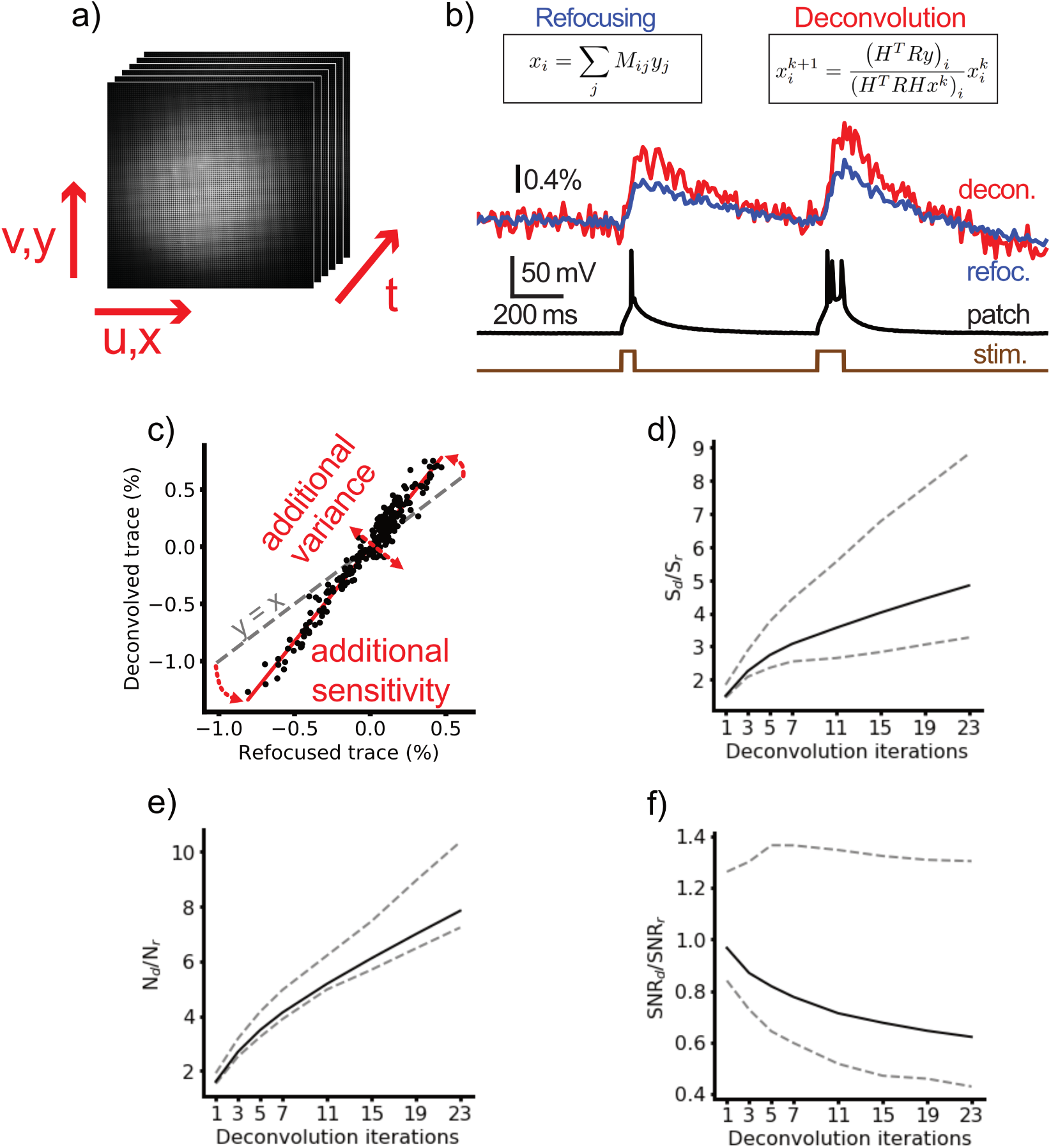
Comparison of different reconstruction methods on SNR. a) Light field time series were collected of functional voltage signals from sparsely expressed GEVIs. b) Time series were extracted from in focus image sequences of the soma via refocusing (left) and ISRA deconvolution (right) and the signal and noise were compared. c) Deconvolved and refocused signals are strongly linearly correlated, as can be seen from plotting the individual trace time points. The additional noise variance due to deconvolution can be identified as the residual from the linear fit. The increased signal level can be seen as the increased fit gradient over unit slope (grey dashed line). Both the noise d) and the signal e) increase monotonically with increasing deconvolution iteration, leading to an overall reduction in SNR with iteration number f). At low iteration number the deconvolution and refocusing are very similar. At large iteration number the SNR is decreased relative to refocused; however, increased axial sectioning may still motivate the use of deconvolution methods. Solid lines are median of n = 15 cells, dashed lines indicate 25^th^ and 75^th^ percentile values. Traces in b) were generated from an average of 8 sweeps.

**Fig 3.**
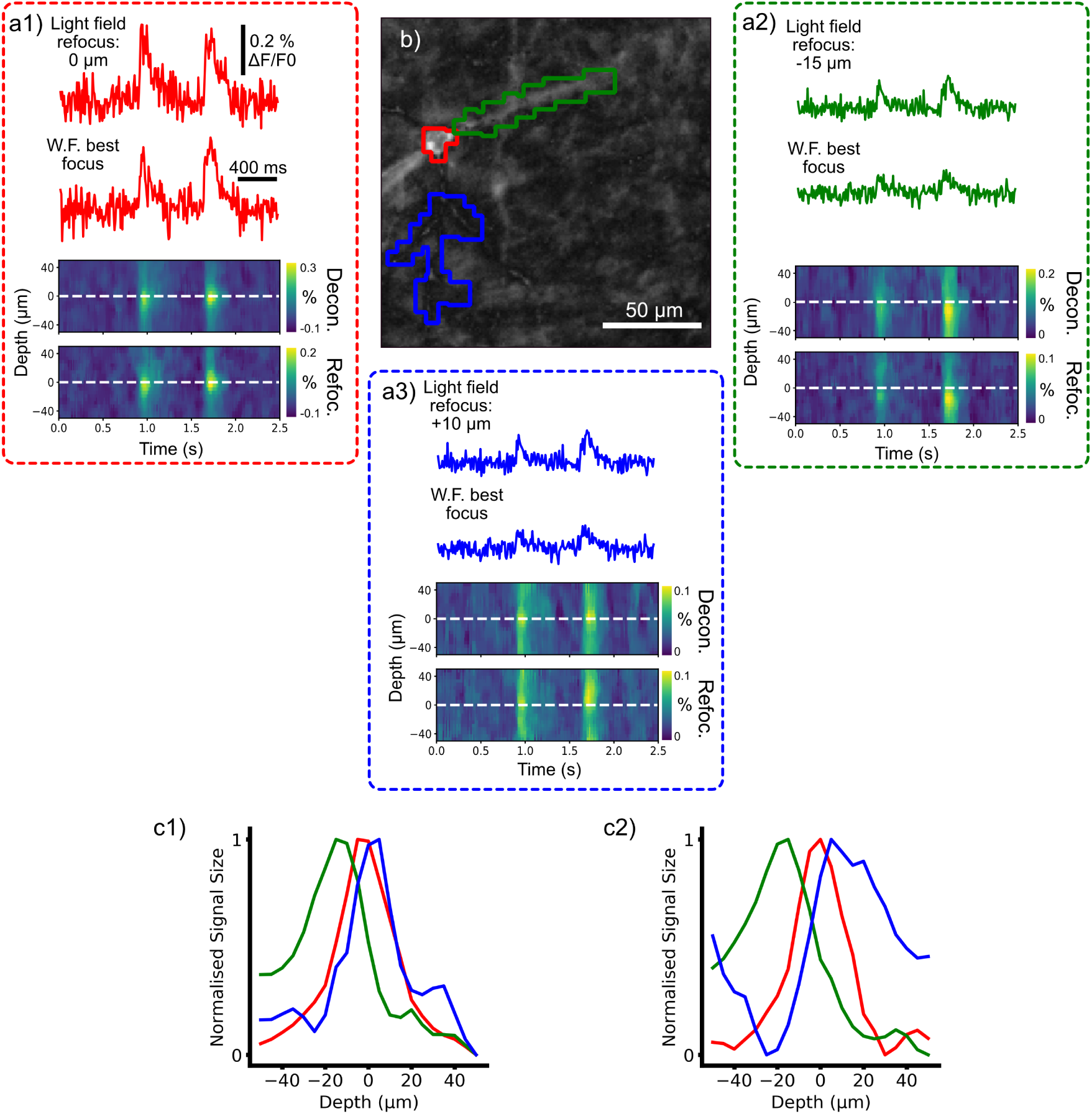
Deconvolution Light field microscopy resolves 3D localised voltage signals. a1-3) Time courses and depth-time plots showing signals from different cellular compartments (shown in (b)) localised at different depths. The somatic signal (a1) is maximal in the wide field and native light field focal planes, whilst the apical dendrite (a2) descends into the slice with its ROI localised 15 um deeper. The signal from a basal dendrite (a3) is superficial to the soma, and its best focal depth is difficult to localize due to the broad axial extent of the refocused signal. The basal and apical dendritic fluorescence transients in the wide field time courses have smaller signals than the light field signals as they are out of plane when focused on the soma. c) The normalised signal size for each ROI across different deconvolved (c1) and refocused (c2) depths. Deconvolution increases the axial localisation of signals. The data are an average of 8 sweeps.

#### 2.3.1 Depth-time plots

To determine the center of mass of the signal for different cellular ROIs we extracted time courses from each refocused or deconvolved depth, then filtered the resulting depth-time 2D arrays with a median filter of 11 samples in the time axis (110 ms) and 3 samples in the z axis (15 µm).

#### 2.3.3 Comparison of light field and wide field SNR

We compared the SNR between trials of the same cell for image sequences taken with wide field and light field imaging systems. We compared the SNR between refocused and wide field images for the same number of repeats using ROIs calculated to be the same for both imaging modalities. For 8/12 cells, an extra aperture was introduced into our light field microscope to compensate for chromatic aberration, reducing the light throughput of the microscope by between 1/2 and 3/4 during light field imaging compared to the equivalent wide field trials. To account for this, the SNR for these trials was adjusted by a factor equal to the square root of the ratio of the mean brightness of the first imaging trials from the light field and wide field trials. The microscope was realigned to account for chromatic aberration before the final 4/12 cells, meaning the design light throughput of the microscope was the same between light field and wide field trials. For these trials the raw SNR was included in the analysis.

#### 2.3.4 Signal spread analysis

We compared the lateral and axial signal spread using a method similar to our previous work.^65^ We quantified the neuronal voltage signal strength in each pixel to create 2- or 3-D ‘activation maps’ by calculating the temporal correlation coefficient of each pixel’s time course with a seed time course from the somatic ROI.

We compared the spatial autocorrelations of these activation maps to quantify the average signal crosstalk between cellular voltage signals.^65^ In our previous work we described how the auto-correlation can be used to quantify the average signal power a cell contributes to a specific pixel’s time course and quantified this effect for this preparation with wide field microscopy in 2 dimensions.^65^ In this work, we calculated the 3D autocorrelations of the lightfield volumes and 2D autocorrelations of the wide field volumes using Fast Fourier Transform based convolution, setting the central 10 *×* 10 pixels of the autocorrelations to the mean of their perimeter to remove a central noise peak.

## 3 Results

### 3.1 Light Field Microscopy Enables Simultaneous Imaging of Axially Separated Dendrites

We demonstrated LFM’s ability to resolve axially separated structures by imaging a cell with a complex 3D dendritic arbour using both wide field and light field microscopy (Fig. 1b) and LFM (Fig. 1c). No single plane wide field image was able to simultaneously bring all the dendrites into a good focus (Fig. 1d), however in different planes from a volume reconstructed by deconvolution different dendritic structures could be clearly distinguished (Fig. 1e1 & Fig. 1e2). The same cellular features can clearly be seen in a standard deviation projection through the reconstructed LFM stack (Fig. 1e3) and a wide field z stack through the same cell (Fig. 1b, both projections through stacks at 1 µm axial increments).

### 3.2 Comparison of the Effect of Different Reconstruction Methods on Signal-to-Noise Ratio

Low-sensitivity GEVIs mean SNR is of utmost importance in voltage imaging analysis strategies, and so we first compared the performance of the deconvolution and refocusing reconstruction approaches on this metric. We reconstructed volume time series for 15 cells from LF time series (Fig 2a) and extracted optical time courses from ROIs over the individual cell’s soma at the native focal plane and compared the SNR between deconvolved and refocused volumes (Fig. 2b). Commonly used LFM iterative reconstruction schemes are prone to noise amplification^78^ which increases with iteration number. It is therefore crucial to understand when to stop the iteration scheme. We used the refocused images as a baseline comparison for the iteration analysis due to the ease of their reconstruction. We found that for all iteration numbers the noise and signal level was increased by deconvolution which increased sensitivity and variance (Figs. 2c - e). The signal significantly increased from 0.3 (0.2, 0.4) % (all results presented as median (IQR)) in the refocused time series to 1.4 (0.9, 1.7) % for the 21 iteration deconvolved traces (Wilcoxon signed rank, *n* = 15, *z* = 0.0, *p* = 0.0003). The noise significantly increased from 0.05 (0.04, 0.08) % in the refocused time series to 0.4 (0.3, 0.5) % for the 21 iteration deconvolved traces (Wilcoxon signed rank, *n* = 15, *z* = 0.0, *p* = 0.0002). This resulted in the SNR reducing from approximately the same as the refocused (1.0 (0.8,1.3)) case for a single deconvolution iteration to around half that of the refocused case (0.6, (0.4, 1.3)). We also processed the light field time series using RL deconvolution and found no substantial differences compared to ISRA.

### 3.3 Light Field Microscopy Resolves 3D Localised and Axially Separated Voltage Signals

We then explored a key advantage of sub-cellular resolution light field voltage imaging: 3D imaging of neuronal processes. Achieving this requires signals from different planes to be discriminable in volume reconstructions. Axial discriminability depends on intrinsic factors such as depth of field, and also extrinsic factors, such as cellular morphology and signal spread due to tissue scattering. To demonstrate the resolution of subcellular voltage transients in 3D, we reconstructed 4D (x,y,z,t) volumes from light field image time series and compared the temporal signals from ROIs over different dendritic and somatic structures in multiple axial planes.

Figure 3 demonstrates LF imaging’s ability to axially localize functional voltage signals from neuronal processes and thereby image functional activity in 3D with SNR unachievable by any equivalent wide field system. Panels 3a 1-3) show single plane and multi-plane time courses from three different ROIs over cellular compartments from a neuron distributed over multiple axial planes. A somatic ROI (Fig. 3a1) in the native focal plane of both wide field and light field images contains action potential evoked fluorescence transients approximately equal in signal size for both light field and wide field (top). The depth-time plots show the functional signal localised to the native focal plane in the LF functional stacks (bottom). In contrast, an ROI over the apical dendrite (Fig. 3(1) has the largest signal when the LF image is refocused 15 µm deeper into the slice, and the signal in the equivalent wide field ROI is much smaller. The depth-time plots for this ROI from both deconvolved and refocused stacks also clearly show the center of mass of the signal located deeper than the native focal plane (Fig. 3(1), bottom). Signals from a basal dendrite (Fig. 3(3) are similarly larger in the LF image refocused 10 µm shallower than the native focal plane. The corresponding depth-time plots show a slight shift in the signal center of mass to a shallower depth, especially in the refocused case.

Plots of signal size as a function of depth for the refocused and deconvolved cases (Fig. 3c 1 & 2) show the axial localization as distinctly different planes for each ROI and also demonstrate a key advantage of deconvolved over refocused reconstructions: increased accuracy in axial localization of functional signals.

### 3.4 Deconvolution Increases Axial Localisation of Functional Voltage Signals

The transients from refocused volumes exhibit a larger axial PSF width compared to the deconvolved traces (Fig. 3c). Hence these signals are smeared out, reducing distinguishability of signal contributions from different planes. To quantify this effect we generated volumes showing the distribution of functional signal. We generated a time course from an in focus somatic ROI and calculated the temporal correlation coefficient of every pixel in the volume for refocused and deconvolved volume time series. Pixels with high correlation coefficients are interpreted as having a large response to the intracellular current injection, and so a volume map of these reveals morphology of structures through which the functional signal propagates. Activation maps from wide field imaging trials show blurring around the soma from out of focus basal dendrites (Fig. 4a). Comparatively, *z*-projections from a 70 µm region around the soma generated from the deconvolved activation volume (Fig. 4b) reveals the structures that cause this blur. A projection through 70 µm around the focus from the refocused case shows significantly more blurring due to the poor axial sectioning of this technique (Fig. 4c). We used the 3D autocorrelation of these activation maps to quantify the spread of the signal in 3D (see section 2.3.4). We quantified how the peak autocorrelation from each cell, and therefore functional signal contribution, decayed axially. Axial smearing can be seen in reconstructions from both deconvolution and refocusing (Fig. 4d), although the effect is much more severe in refocused traces. The smearing appears in the axial autocorrelation as both broadening of the central peak and increased side lobes (Fig. 4e). The central peak width and side lobes decrease with increased deconvolution iteration number (Fig. 4f & g), thus increasing the axial sectioning.

**Fig 4.**
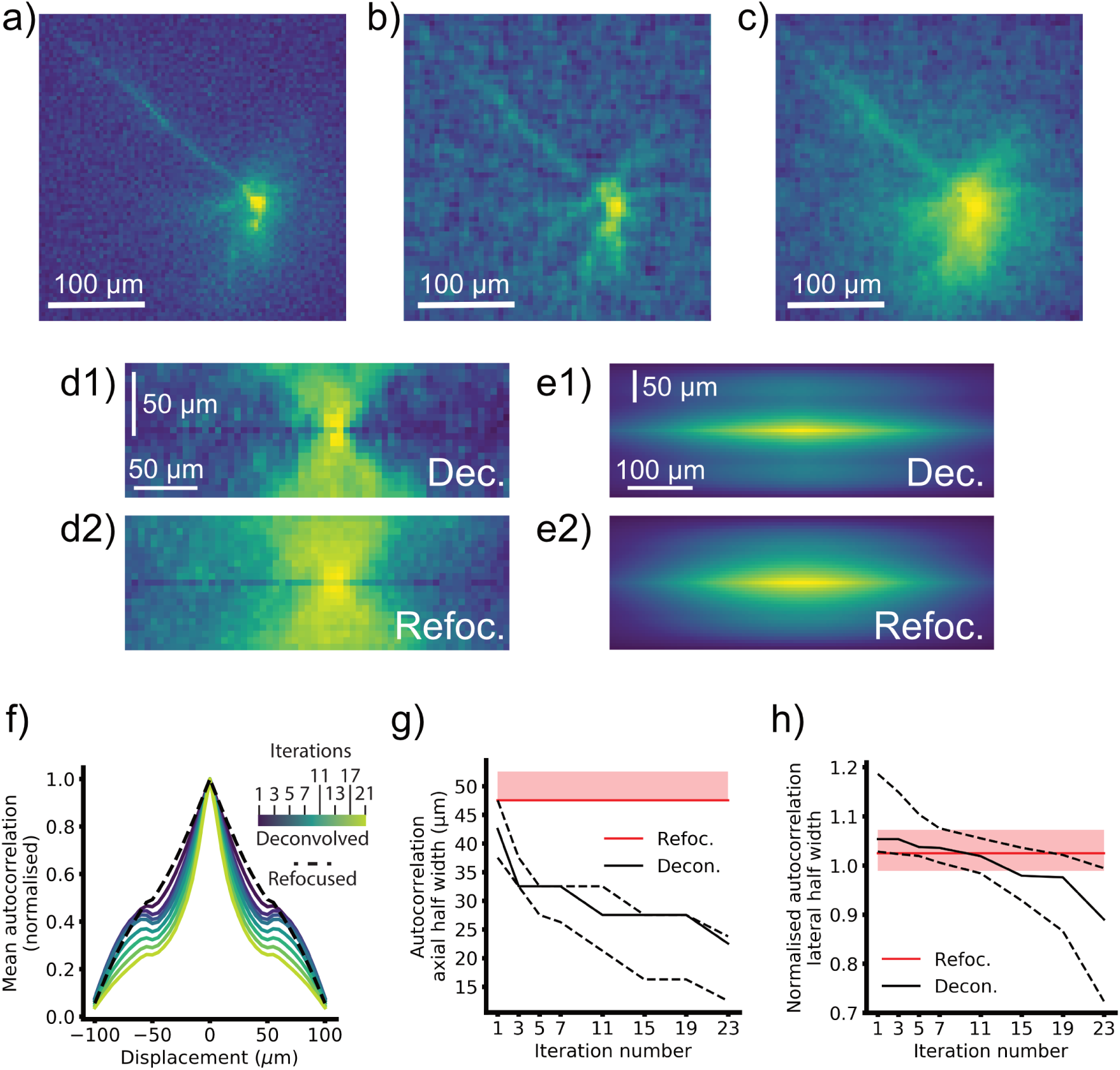
Mapping dendritic signals. a) wide field activation image. b) Deconvolved activation image, sum projection from −35 to +35 um, 5 deconvolution iterations. c) Refocused activation image, sum projection from −35 to +35 um. d) x-z maximum intensity projections through deconvolved (top), and refocused (bottom) activation images showing the different axial sectioning. e) Mean x-z projections through the autocorrelations. f) Normalised maximum autocorrelation for different depths from refocused and deconvolved LFM activation volumes. The secondary peaks arise from the elongated axial PSF, and these can be seen decreasing as the iteration number increases. g) Median autocorrelation axial half widths for n = 12 cells with iteration number. Dashed lines represent quartile values. Red line is the refocused median width and shaded area the refocused interquartile range (IQR). h) Median autocorrelation lateral widths normalised to wide field lateral widths for refocused images (red, shaded area IQR) and different deconvolution iterations (black lines, dashed lines IQR).

**Fig 5.**
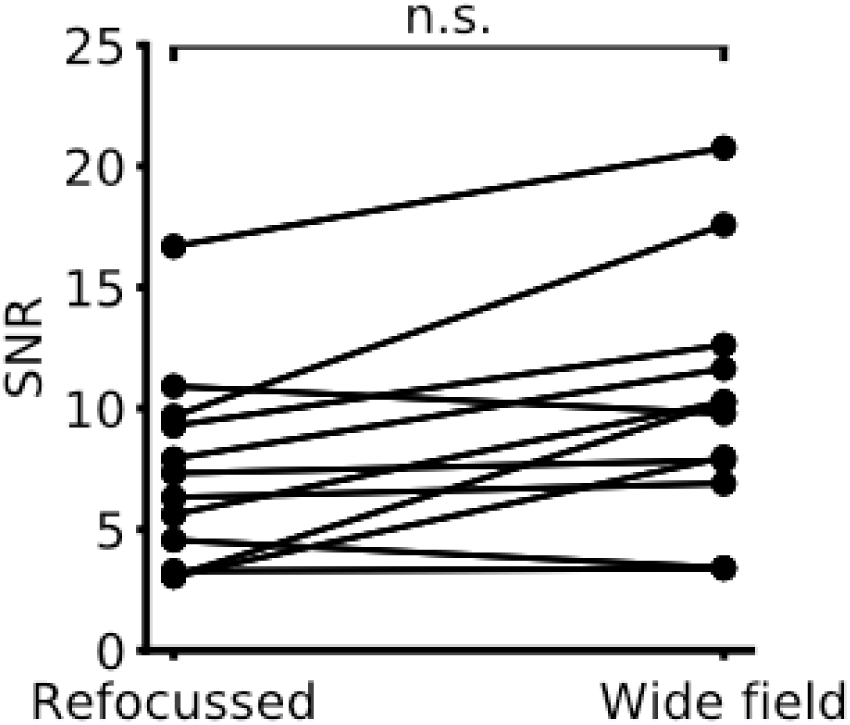
Comparison of Light field and Wide Field SNR. Points correspond to mean SNR between paired light field and wide field trials. Light field microscopy SNR does not differ significantly from wide field. For 8/12 trials we included a correction factor due to a misalignment in the LFM as discussed in section 2.3.3.

The autocorrelation widths decreased significantly from 1 to 21 iterations (median dropped from 42.5 (37.5, 47.5) µm to 22.5 (12.5, 23.75) µm, *z* = 0, *p* = 0.002), and for both cases the axial spread was significantly lower than refocused (median of 47.5 (47.5, 52.5) µm, *p* = 0.001, *p* = 0.002 for 1 and 21 iterations respectively). Significance tests were performed with a Friedman *χ*^2^ with post hoc Bonferroni-corrected Wilcoxon signed-rank tests (significant at *p <* 0.017). n = 12 cells from 12 slices from 4 mice. Friedman *χ*^2^ = 24, *p* = 6 *×* 10^*−*6^.

Finally we compared how deconvolution and refocusing affected lateral signal localization compared to the equivalent wide field time series (Fig. 4h). We measured the width of radially averaged autocorrelations normalised to matched wide field trials for the refocused and deconvolved cases. We found that the lateral signal spread significantly decreased from 1 to 21 iterations (median dropped from 1.05 (1.03, 1.18) times larger than wide field trials to 0.89 (0.73, 0.99) times larger, *Z* = 0, *p* = 0.002), from significantly larger than the WF and refocused at 1 iteration (*z* = 1, *p* = 0.003 and *z* = 0, *p* = 0.002 respectively), to significantly smaller than the refocused at 21 iterations (*z* = 0.0, *p* = 0.003). The refocused widths did not differ significantly from the matched wide field trials (median 1.02 (0.99, 1.07) times larger, *z* = 17, *p* = 0.08). Significance tests were performed with a Friedman *χ*^2^ with post hoc Bonferroni-corrected Wilcoxon signedrank tests on the raw widths (significant at *p <* 0.0083). *n* = 12 cells from 12 slices from 4 mice. Friedman *χ*^2^ = 26, *p* = 9 *×* 10^*−*6^.

In total our analyses reveal that deconvolution improves both axial and lateral signal localization, but decreases temporal signal SNR compared to synthetic refocussing, with both effects intensifying with increasing iteration number.

### 3.5 Temporal signal SNR is Unaffected by Light Field Imaging

We measured the SNR for paired wide field and refocused light field imaging trials in the same cells. For 8/12 trials we included a correction factor due to a misalignment in the LFM as discussed in section 2.3.3. The SNR did not change significantly between the light field and wide field cases (Wilcoxon signed rank test, n = 12 cells from 12 slices and 4 mice, *z* = 32, *p* = 0.6), with a median light field SNR of 8.4 (5.2, 11.4) and a median wide field SNR of 10.0 (7.6, 11.9).

## 4 Discussion

We have shown that LFM enables 3D sub-cellular GEVI imaging of somatic and dendritic structures. We demonstrated that LFM enables simultaneous imaging of axially separated dendrites, overcoming a key limitation of wide field imaging. We further showed that functional voltage signals from dendrites could be axially resolved at different depths. This finding is key to demonstrating LFM’s utility for studies of dendritic integration or synaptic mapping.

We compared how synthetic refocussing and deconvolution-based reconstruction techniques perform with respect to spatial signal localization and temporal SNR. Synthetic refocussing is computationally simple and can be used to process light fields online, during an experiment, or post hoc. Refocusing features better temporal signal SNR but poorer lateral and axial confinement compared to deconvolution. Deconvolution has two major disadvantages: computational cost and noise amplification. As the light field microscope PSF is not shift invariant it is described by a 5 dimensional matrix, complicating reconstruction. The periodicity it displays under lateral shifts by integer multiples of the microlens pitch, however, enable deconvolution to be performed efficiently by using FFT-based convolutions. Despite this, even small increases in lateral sampling in the deconvolved volume increase the computational cost of reconstruction drastically. Reconstructing *n*_*z*_ *z*-planes in a volume with a lateral increase in sampling over the native LFM sampling of *m* requires 2 *× n*_*z*_ *× m*^2^ 2D convolutions per iteration, precluding online image processing. Secondly, both Richardson-Lucy and ISRA tend to amplify noise in their outputs due to their lack of regularization.^78^ This noise may be acceptable when imaging high-SNR calcium signals, however it can dominate small, dim voltage signals. Incorporating regularization into the deconvolution approaches to suppress noise overfitting could also ameliorate deconvolution’s effects on temporal SNR.

In this study we imaged VSFP-Butterfly 1.2, an older generation probe. GEVI technology has advanced dramatically recently, greatly increasing their sensitivity, and with these new sensors noise amplification due to deconvolution in the light field volume reconstruction may become less significant. Although VSFP-Butterfly 1.2 exhibits lower sensitivity than several recently reported probes,^20, 79–83^ we were able to express it sparsely and strongly to enable single-cell GEVI imaging without somatic restriction, which would preclude study of sub-cellular signals.^65, 66^ The slow kinetics of the probe used in this study also enabled resolution of action potentials at 100 frames/s without severe aliasing. Although we could resolve single-sweep signals, signal averaging was required to resolve smaller dendritic signals with adequate SNR. With a more recent GEVI, dendritic processes could likely be resolved in single sweeps.

Newer voltage sensors can not immediately be combined with LFM, however, as they require much faster sampling rates, typically between 500 - 1000 Hz. Megapixel cameras with 1 kHz full-frame readout rates are therefore needed to fully exploit these newer voltage indicators. Current sCMOS cameras such as the one used in this study can achieve these imaging rates by reducing the FOV to a small central strip of the image sensor. This, however, is particularly detrimental to LFM compared to wide field imaging as the LFM PSF spreads information about each point widely across the image sensor for objects away from the focal plane. If only a small strip of the sensor is imaged SNR will be greatly degraded as light is lost outside of this reduced FOV. We anticipate that this issue will be steadily ameliorated as faster sCMOS sensor technology is developed.

A second issue arises with newer, faster GEVIs due to their requirement for much faster frame rates. Deconvolving individual frames with these sensors would require a drastic increase in computational resources and is likely untenable. Approaches have been developed for calcium imaging light field time series which do not involve deconvolution of every frame.^61, 84^ In their current form, however, these are unsuitable for reconstruction of subcellular light field voltage imaging time series as they leverage the temporal and spatial characteristics of neuronal calcium imaging as reconstruction priors. These priors, such as somatic signal localisation or sparse temporal activity, are not as applicable to subcellular voltage imaging signals, which are smaller, less temporally sparse and arise from more morphologically intricate structures than neuronal somata.

Finally, in this study we compared the SNR between refocused LFM volumes and matched wide field traces and found they did not differ significantly. This is expected, as apart from light losses at the MLA, which are *<* 15% according to the manufacturer, there are no significant losses of SNR to shot noise between wide field and light field microscopy. Together these results have the potential to motivate further work and widespread application of light field microscopy to voltage imaging owing to light fields high photon budget and ability to resolve neurons in three spatial dimensions.

## Acknowledgements

We thank Yu Liu for her technical assistance and support for research in the Foust and Schultz labs. The authors would also like to thank the Imperial College Research Computing Service.

This work was supported by the following grants: Engineering and Physical Sciences Research Council (EP/L016737/1); Wellcome Trust Seed Award (201964/Z/16/Z); The Royal Academy of Engineering under the RAEng Research Fellowships scheme (RF1415/14/26); Biotechnology and Biology Research Council (BB/R009007/1 and BB/R022437/1); The Royal Society (TA/R1/170047); The BRAIN initiative (US National Institutes of Health, U01MH109091, U01NS099573).

## Author Contributions

PQ, CLH, SRS and AJF conceived and designed the experiments. CLH and AJF designed the light field optics. PQ and CLH performed experiments. PQ, CLH, AJF and SRS designed the analysis. PQ, HVJ, PS, and PLD developed the deconvolution approach. CS and TK developed and prepared animals with strong GEVI expression limited to a sparse neuronal subpopulation. PQ and MN developed the light field PSF model. PQ analyzed the data and wrote the paper. All authors contributed to manuscript revision and approved the final manuscript.

## Data, Materials, and Code Availability

The datasets and code generated for this study are available on request to the corresponding author.

